## Supplementary file for "AI in Practice: A Multilingual Survey of 2025 BioHackathon Participants"

### Biohackathon 2025 AI Survey

*EN:*

We want to learn how Biohackathon participants are using AI in their work.

Please share your experiences—the tools you use, successes you've had, and challenges you've faced.

If you are actively applying AI in your research, we invite you to give a brief review of your work so it can be cited and highlighted. Your responses will help build a community resource and will be shared back with all participants.

**Definition of AI (for this survey)**

Here, "*AI*" includes machine learning, deep learning, and generative AI tools used in research and professional work. Examples:

- Large Language Models
- Scientific AI tools
- Machine learning libraries or custom-trained models
- AI-powered tools for coding, data analysis, or literature review

Traditional statistical methods are not included unless they involve AI/ML.
The survey should take 10-15 minutes to complete!

*日本語:*
バイオハッカソン参加者の皆さまへ

本調査では、参加者の皆さまが業務や研究においてAIをどのように活用されているかをお伺いします。

ご自身のご経験（使用されているツール、成功事例、直面した課題など）についてぜひお聞かせください。

研究でAIを積極的にご活用されている方は、研究の概要を簡潔にご紹介いただければ幸いです。ご紹介いただいた内容は引用や事例紹介として取り上げさせていただく場合があります。皆さまのご回答はコミュニティリソースの構築に役立てられ、全参加者と共有されます。

本調査における「**AI**」の定義

本調査における「AI」には、研究や専門的な業務で利用される機械学習、深層学習、生成AIツールなどが含まれます。例としては以下の通りです。

- 大規模言語モデル
- 科学研究向けAIツール
- 機械学習ライブラリや独自に学習させたモデル
- コーディング、データ分析、文献調査に活用するAI搭載ツール

なお、AI/MLを用いない従来型の統計手法は対象外です。

所要時間

回答に必要な時間はおよそ10〜15分です。

ご協力のほどよろしくお願いいたします。

ภาษาไทย:

มาร่วมแชร์ประสบการณ์การใช้ AI ในงานวิจัยของคุณกัน

เราอยากเรียนรู้ว่าผู้เข้าร่วม Biohackathon นำปัญญาประดิษฐ์ (AI) มาปรับใช้ในการทำงานกันอย่างไรบ้าง

ขอเชิญชวนทุกท่านมาร่วมแบ่งปันประสบการณ์ ไม่ว่าจะเป็นเครื่องมือที่ใช้ ความสำเร็จ หรือความท้าทายที่เคยเจอ

หากคุณกำลังใช้ AI ในงานวิจัยของคุณอยู่ ขอเชิญร่วมรีวิวสั้นๆ เกี่ยวกับงานของคุณ เพื่อที่เราจะสามารถนำไปอ้างอิงและนำเสนอ คำตอบของคุณจะเป็นส่วนสำคัญในการสร้างแหล่งข้อมูลสำหรับชุมชน และเราจะนำข้อมูลเหล่านี้กลับมาแบ่งปันให้ผู้เข้าร่วมทุกคนได้ใช้ประโยชน์ต่อไป

**คำนิยามของ AI ในแบบสำรวจนี้**

ในที่นี้ AI หมายถึง เทคโนโลยีการเรียนรู้ของเครื่อง (Machine Learning) การเรียนรู้เชิงลึก (Deep Learning) และเครื่องมือ AI ประเภท Generative AI ที่ใช้ในงานวิจัยและการทำงาน ตัวอย่างเช่น:

- แบบจำลองภาษาขนาดใหญ่ (Large Language Models)
- เครื่องมือ AI ทางวิทยาศาสตร์โดยเฉพาะ
- ไลบรารี Machine Learning หรือโมเดลที่สร้างและฝึกฝนขึ้นเอง
- เครื่องมือที่ขับเคลื่อนด้วย AI สำหรับการเขียนโค้ด การวิเคราะห์ข้อมูล หรือการทบทวนวรรณกรรมทางวิชาการ

ทั้งนี้ AI จะไม่รวมวิธีการทางสถิติแบบดั้งเดิม ยกเว้นในกรณีที่มีการใช้ AI/ML เข้ามาเกี่ยวข้อง

แบบสำรวจนี้ใช้เวลาประมาณ 10 - 15 นาที

* Indicates required question

Please choose your language / 言語を選んでください / โปรดเลือกภาษาสำหรับแบบสอบถาม

1. Select language *

*Mark only one oval.*

- ⬭ 日本語 *Skip to question 46*
- ⬭ English *Skip to question 2*
- ⬭ ภาษาไทย *Skip to question 24*

Demographic Information

Please tell us a little about yourself!

2. Are/were you a BioHackathon participant? (Select all that apply)

*Check all that apply.*

- ☐ 2025 BioHackathon participant in Japan
- ☐ Participant of previous BioHackathons (any country)
- ☐ Not a BioHackathon participant

3. What is your field? (Select all the apply)

*Check all that apply.*

- [ ] Sequence analysis
- [ ] Genomics and transcriptomics
- [ ] Metagenomics and microbiome informatics
- [ ] Epigenomics and regulatory genomics
- [ ] Proteomics and metabolomics
- [ ] Structural bioinformatics
- [ ] Systems biology
- [ ] Computational evolutionary biology
- [ ] Cheminformatics
- [ ] Medical/clinical informatics
- [ ] Biostatistics
- [ ] Machine learning and AI
- [ ] Data visualization
- [ ] Data standards and interoperability
- [ ] Ontology creation
- [ ] Database management
- [ ] HPC and cloud bioinformatics
- [ ] Front-end development
- [ ] Back-end development
- [ ] Other: ______________________

4. What institution type do you work for?

*Check all that apply.*

- [ ] Academia
- [ ] Public sector
- [ ] Private sector
- [ ] Other: ______________________

5. Which country do you work in?

______________________

6. What is your age?

*Mark only one oval.*

- < 25 years old
- 25 - 34
- 35 - 44
- 45 - 54
- > 55 years old
- Prefer not to answer

7. Which of the following best describes your gender?

*Mark only one oval.*

- Woman
- Man
- Non-binary
- Prefer not to answer
- Other: ______________________

8. Which of the following best describes your use of AI? *

*Mark only one oval.*

- I am an AI researcher *Skip to question 10*
- I am an AI developer *Skip to question 10*
- I use AI everyday *Skip to question 12*
- I use AI a few times a week *Skip to question 12*
- I use AI a few times a month *Skip to question 12*
- I never use AI *Skip to question 9*
- Other: ______________________

You don't use AI...

9. Is there a reason why you don't use AI? (Select all that apply)

*Check all that apply.*

- [ ] I don't trust the companies who make AI.
- [ ] AI as a technology is not useful for my work.
- [ ] It's too expensive.
- [ ] I don't trust the results or underlying technology.
- [ ] Other: ______________________

*Skip to question 22*

AI Researcher or Developer

10. Please briefly describe your AI-related work

_____________________________________________
_____________________________________________
_____________________________________________
_____________________________________________
_____________________________________________

11. If you would like to have your work cited please provide us a link to a paper, code repository, etc.

_____________________________________________
_____________________________________________
_____________________________________________
_____________________________________________
_____________________________________________

*Skip to question 12*

Applications and Impact

12. Which AI tools do you use most often?

Separate your answers with ","

______________________________________________

______________________________________________

______________________________________________

______________________________________________

______________________________________________

13. What tasks do you use AI for? When you use AI for that task, which of the following best describes your approach?

*Check all that apply.*

| | I use AI to complete the entire task with little to no editing | I use AI to generate a first draft, which I then edit and finalize | I use AI to assist with specific parts of a task |
|---|---|---|---|
| **Coding** | ☐ | ☐ | ☐ |
| **Research** | ☐ | ☐ | ☐ |
| **Brainstorming** | ☐ | ☐ | ☐ |
| **Writing/Editing** | ☐ | ☐ | ☐ |
| **Teaching and curriculum** | ☐ | ☐ | ☐ |
| **Translation** | ☐ | ☐ | ☐ |
| **Personal use (trip planning, recipes, making songs, etc)** | ☐ | ☐ | ☐ |

14. What has been your success using AI in your work or projects? Please describe the task, the AI tool(s) used, and the outcome.

15. When using AI tools in your work, what challenges have you faced? (Select all that apply)

*Check all that apply.*

- [ ] AI tools are too complex or not user-friendly
- [ ] AI outputs are inaccurate or unreliable
- [ ] AI suggestions are difficult to interpret
- [ ] Difficulty integrating AI into existing workflows
- [ ] Lack of institutional support (software, hardware, guidance)
- [ ] High computational cost / resources required
- [ ] Data limitations (size, quality, accessibility)
- [ ] Ethical or privacy concerns
- [ ] Over-reliance on AI leading to mistakes
- [ ] Other: ______

16. Tell us about a time AI really failed you or caused unexpected trouble. What happened, and what did you learn?

17. Please rate your overall satisfaction with your use of AI.

*Mark only one oval.*

- ( ) 1 (Not at all) *Skip to question 18*
- ( ) 2 *Skip to question 18*
- ( ) 3 *Skip to question 18*
- ( ) 4 *Skip to question 19*
- ( ) 5 (Very satisfied) *Skip to question 19*

*Skip to question 18*

#### AI improvement suggestion(s)

18. What would AI tools need to improve for you to be more satisfied? (Select all the apply)

*Check all that apply.*

- [ ] Accuracy & reliability
- [ ] Usability & efficiency
- [ ] Control & customization
- [ ] Privacy, security & ethics
- [ ] Other: ______________________

*Skip to question 19*

#### Institutional Support

19. How would you describe the level of institutional support you receive regarding AI usage?

*Mark only one oval.*

| | 1 | 2 | 3 | 4 | 5 | |
|---|---|---|---|---|---|---|
| No i | ( ) | ( ) | ( ) | ( ) | ( ) | Very well supported |

20. What kinds of support does your institution provide? Select as many as apply

*Check all that apply.*

- [ ] Provides its own self-hosted AI services
- [ ] Access to institutional AI software or platforms (licenses, subscriptions)
- [ ] Access to high-performance computing / GPU resources
- [ ] Dedicated AI support staff or help desk
- [ ] Training/workshops on AI tools or prompt engineering
- [ ] Support for integrating AI into research workflows
- [ ] Guidance on ethical use of AI / AI governance policies
- [ ] Funding or grants for AI-related projects
- [ ] Other: ____________________

*Skip to question 21*

#### Ethics and Responsibility

21. How concerned are you with the following issues in relation to AI?

*Mark only one oval per row.*

| | 1 - Not concerned at all | 2 | 3 | 4 | 5 - Very concerned |
|---|---|---|---|---|---|
| **Bias in algorithms** | ◯ | ◯ | ◯ | ◯ | ◯ |
| **Data privacy/security** | ◯ | ◯ | ◯ | ◯ | ◯ |
| **Intellectual property, ownership** | ◯ | ◯ | ◯ | ◯ | ◯ |
| **Misinformation/Hallucinations** | ◯ | ◯ | ◯ | ◯ | ◯ |
| **Environment impact** | ◯ | ◯ | ◯ | ◯ | ◯ |

*Skip to question 22*

#### Final thoughts

22. Is there anything else on the topic you'd like to share? Anything we didn't ask but should have?

23. If you're ok with us contacting you with further questions, please share your email here

**ข้อมูลทั่วไป**

กรุณาให้ข้อมูลเกี่ยวกับตัวคุณ

24. คุณเคยเข้าร่วมงานBioHackathon หรือไม่ (เลือกได้มากกว่า 1 ข้อ)

*Check all that apply.*

- [ ] เป็นผู้เข้าร่วม 2025 BioHackathon ที่ประเทศญี่ปุ่น
- [ ] เคยเข้าร่วม BioHackathon ครั้งก่อนๆ (ในประเทศใดก็ได้)
- [ ] ไม่เคยเข้าร่วม BioHackathon

25. โปรดเลือกสายงานวิจัยของท่าน (เลือกได้มากกว่า 1 ข้อ)

*Check all that apply.*

- [ ] การวิเคราะห์ลำดับพันธุกรรม (Sequence analysis)
- [ ] จีโนมิกส์และทรานสคริปโตมิกส์ (Genomics and transcriptomics)
- [ ] เมตาจีโนมิกส์และไมโครไบโอม (Metagenomics and microbiome informatics)
- [ ] จีโนมิกส์เหนือพันธุกรรมและจีโนมิกส์เชิงกำกับ (Epigenomics and regulatory genomics)
- [ ] โปรตีโอมิกส์และเมแทบอโลมิกส์ (Proteomics and metabolomics)
- [ ] ชีวสารสนเทศศาสตร์โครงสร้าง (Structural bioinformatics)
- [ ] ชีววิทยาระบบ (Systems biology)
- [ ] ชีววิทยาเชิงวิวัฒนาการเชิงคำนวณ (Computational evolutionary biology)
- [ ] เคมีสารสนเทศ (Cheminformatics)
- [ ] สารสนเทศทางการแพทย์/คลินิก (Medical/clinical informatics)
- [ ] ชีวสถิติ (Biostatistics)
- [ ] การเรียนรู้ของคอมพิวเตอร์ และปัญญาประดิษฐ์ (Machine learning and AI)
- [ ] การสร้างภาพข้อมูล (Data visualization)
- [ ] การสร้างออนโทโลยี (Ontology creation)
- [ ] มาตรฐานข้อมูลและการทำงานร่วมกันของข้อมูล (Data standards and interoperability)
- [ ] การจัดการฐานข้อมูล (Database management)
- [ ] ชีวสารสนเทศศาสตร์บนคลาวด์และคอมพิวเตอร์สมรรถนะสูง (HPC and cloud bioinformatics)
- [ ] การพัฒนาส่วนหน้า (Front-end development)
- [ ] การพัฒนาส่วนหลัง (Back-end development)
- [ ] Other: ______________________

26. ท่านปฏิบัติงานอยู่ที่หน่วยงานประเภทใด

*Check all that apply.*

- [ ] มหาวิทยาลัย
- [ ] หน่วยงานภาครัฐ
- [ ] หน่วยงานภาคเอกชน
- [ ] Other: ______________________

27. ท่านประกอบอาชีพอยู่ที่ประเทศใด

______________________

28. โปรดเลือกช่วงอายุปัจจุบันของท่าน

*Mark only one oval.*

- < 25 ปี
- 25 - 34 ปี
- 35 - 44 ปี
- 45 - 54 ปี
- 55+ ปี
- ไม่สะดวกให้ข้อมูล

29. ข้อใดระบุถึงเพศสภาพของท่านได้ดีที่สุด

*Mark only one oval.*

- ผู้ชาย
- ผู้หญิง
- เพศทางเลือก
- ไม่สะดวกให้ข้อมูล
- Other: ______________________

30. ข้อใดอธิบายการใช้งาน AI ของคุณได้ดีที่สุด *

*Mark only one oval.*

- นักวิจัยเกี่ยวกับ AI (AI researcher) *Skip to question 32*
- นักพัฒนา AI (AI developer) *Skip to question 32*
- ใช้ AI ทุกวัน *Skip to question 34*
- ใช้ AI หลายครั้งในหนึ่งสัปดาห์ *Skip to question 34*
- ใช้ AI หลายครั้งในหนึ่งเดือน *Skip to question 34*
- ไม่เคยใช้ AI *Skip to question 31*
- Other: ______________________

สำหรับท่านไม่เคยใช้ AI

31. อะไรคือเหตุผลที่ท่านไม่ใช้ AI (เลือกได้มากกว่า 1 ข้อ)

*Check all that apply.*

- [ ] ไม่ไว้วางใจบริษัทที่สร้าง AI
- [ ] เทคโนโลยี AI ไม่มีประโยชน์ต่องาน
- [ ] มีค่าใช้จ่ายสูงเกินไป
- [ ] ไม่เชื่อมั่นในผลลัพธ์หรือตัวเทคโนโลยี
- [ ] Other: ____________

*Skip to question 44*

#### สำหรับนักวิจัยเกี่ยวกับ AI และ นักพัฒนา AI

32. โปรดอธิบายงานของท่านที่เกี่ยวข้องกับ AI โดยสังเขป

_____________________________________________
_____________________________________________
_____________________________________________
_____________________________________________
_____________________________________________

33. หากต้องการให้ผลงานของท่านได้รับการอ้างอิง โปรดระบุลิงก์ไปยังผลงาน เช่น บทความวิชาการ, code repository หรืออื่นๆ

_____________________________________________
_____________________________________________
_____________________________________________
_____________________________________________
_____________________________________________

*Skip to question 34*

#### การประยุกต์ใช้และผลกระทบ

34. ท่านใช้เครื่องมือ AI ใดบ้าง (ตอบได้มากกว่า 1 ข้อ)

สำหรับมากกว่า 1 คำตอบ โปรดใช้ "," คั่น

35. ท่านใช้ AI สำหรับงานประเภทใดบ้าง, เมื่อท่านใช้ AI ทำงานด้านนั้น ข้อใดอธิบายวิธีการของคุณได้ดีที่สุด (เลือกได้มากกว่า 1 ข้อ)

*Check all that apply.*

| | ใช้ AI ทำงานทั้งหมดจนเสร็จโดยแก้ไขเพียงเล็กน้อยหรือไม่แก้ไขเลย | ใช้ AI สร้างฉบับร่างแรกจากนั้นจึงนำมาแก้ไขจนสมบูรณ์ | ใช้ AI สำหรับงานบางส่วนเท่านั้น |
|---|---|---|---|
| **การเขียนและแก้โค้ด** | ☐ | ☐ | ☐ |
| **การวิจัย / การค้นคว้าข้อมูล** | ☐ | ☐ | ☐ |
| **การระดมสมอง / การหาไอเดีย** | ☐ | ☐ | ☐ |
| **การเขียน / การแก้ไขข้อความ** | ☐ | ☐ | ☐ |
| **การสอนและหลักสูตร** | ☐ | ☐ | ☐ |
| **การแปลภาษา** | ☐ | ☐ | ☐ |
| การใช้ [illegible] | | | |

| | | | |
|---|---|---|---|
| ...งานส่วนตัว (เช่น วางแผนเที่ยว, หาสูตรอาหาร, แต่งเพลง) / ...สำหรับใช้งานส่วนตัว (เช่น วางแผนเที่ยว, หาสูตรอาหาร, แต่งเพลง) | ☐ | ☐ | ☐ |

36. โปรดเล่าถึงตัวอย่างความสำเร็จครั้งใหญ่ของคุณในการใช้ AI กับงานหรือโปรเจกต์ โดยอธิบายลักษณะของงาน, เครื่องมือ AI ที่ใช้ และผลลัพธ์ที่ได้รับ

________________________________________

________________________________________

________________________________________

________________________________________

________________________________________

37. ท่านพบความท้าทายอะไรบ้างในการใช้เครื่องมือ AI กับการทำงาน (เลือกได้มากกว่า 1 ข้อ)

*Check all that apply.*

- ☐ เครื่องมือ AI ซับซ้อนหรือใช้งานยากเกินไป
- ☐ ผลลัพธ์จาก AI ไม่แม่นยำหรือไม่น่าเชื่อถือ
- ☐ ข้อเสนอแนะจาก AI ตีความหรือนำไปใช้ได้ยาก
- ☐ ปัญหาในการนำ AI มาใช้ร่วมกับขั้นตอนการทำงานที่มีอยู่
- ☐ ขาดการสนับสนุนจากองค์กร (เช่น ซอฟต์แวร์, ฮาร์ดแวร์, คำแนะนำ)
- ☐ ต้นทุนในการประมวลผลสูง / ต้องใช้ทรัพยากรมาก
- ☐ ข้อจำกัดด้านข้อมูล (เช่น ขนาด, คุณภาพ, การเข้าถึง)
- ☐ ข้อกังวลด้านจริยธรรมหรือความเป็นส่วนตัว
- ☐ การพึ่งพา AI มากเกินไปจนนำไปสู่ข้อผิดพลาด

- ☐ Other: ________________________

38. โปรดเล่าถึงเหตุการณ์ที่ AI ทำงานผิดพลาด หรือสร้างปัญหาที่ไม่คาดคิด และได้เรียนรู้อะไรจากประสบการณ์นั้น

39. ความพึงพอใจโดยรวมในการใช้งาน AI

*Mark only one oval.*

- 1 (ไม่พึงพอใจเลย) *Skip to question 40*
- 2 *Skip to question 40*
- 3 *Skip to question 40*
- 4 *Skip to question 41*
- 5 (พึงพอใจอย่างมาก) *Skip to question 41*

#### ข้อเสนอแนะเพื่อการปรับปรุง AI

40. เครื่องมือ AI จำเป็นต้องปรับปรุงในด้านใด ท่านจึงจะพึงพอใจกับการใช้งานมากขึ้น (เลือกได้มากกว่า 1 ข้อ)

*Check all that apply.*

- [ ] ความแม่นยำและความน่าเชื่อถือ
- [ ] การใช้งานง่าย (usability) และประสิทธิภาพ (efficiency)
- [ ] การควบคุมและการปรับแต่ง (control and optimization)
- [ ] ความเป็นส่วนตัว ความปลอดภัย และจริยธรรม
- [ ] Other: ____________

*Skip to question 41*

#### การสนับสนุนจากองค์กร

41. คุณจะอธิบายระดับการสนับสนุนที่คุณได้รับจากองค์กรว่าอย่างไร

*Mark only one oval.*

| | 1 | 2 | 3 | 4 | 5 | |
|---|---|---|---|---|---|---|
| ไม่ไ | ◯ | ◯ | ◯ | ◯ | ◯ | ได้รับการสนับสนุนเป็นอย่างดี |

42. องค์กรของคุณให้การสนับสนุนด้านใดบ้าง (เลือกได้มากกว่า 1 ข้อ)

*Check all that apply.*

- [ ] มีบริการ AI ขององค์กรเอง (self-hosted)
- [ ] การให้สิทธิ์เข้าถึงซอฟต์แวร์หรือแพลตฟอร์ม AI (เช่น license, subscription)
- [ ] การสนับสนุนการเข้าถึงทรัพยากรคอมพิวเตอร์สมรรถนะสูง / GPU
- [ ] มีเจ้าหน้าที่หรือ help desk ที่ให้การสนับสนุนด้าน AI โดยเฉพาะ
- [ ] การจัดฝึกอบรม/เวิร์กช็อปเกี่ยวกับเครื่องมือ AI หรือ prompt engineering
- [ ] การสนับสนุนการนำ AI มาปรับใช้ในขั้นตอนการทำวิจัย
- [ ] คำแนะนำด้านจริยธรรมในการใช้ AI / นโยบายการกำกับดูแล AI
- [ ] เงินทุนสนับสนุนสำหรับโครงการที่เกี่ยวข้องกับ AI
- [ ] Other: ____________________

*Skip to question 43*

#### จริยธรรมและความรับผิดชอบ

43. ท่านมีความกังวลต่อประเด็นต่อไปนี้ที่เกี่ยวข้องกับ AI มากน้อยเพียงใด

*Mark only one oval per row.*

| | 1 - ไม่กังวลเลย | 2 | 3 | 4 | 5 - กังวลมากที่สุด |
|---|---|---|---|---|---|
| **ความลำเอียงในอัลกอริทึม** | ◯ | ◯ | ◯ | ◯ | ◯ |
| **ความเป็นส่วนตัวและความปลอดภัยของข้อมูล** | ◯ | ◯ | ◯ | ◯ | ◯ |
| **ทรัพย์สินทางปัญญาและกรรมสิทธิ์** | ◯ | ◯ | ◯ | ◯ | ◯ |
| **ข้อมูลเท็จและการให้ข้อมูลที่ผิดพลาด (Hallucinations)** | ◯ | ◯ | ◯ | ◯ | ◯ |
| **ผลกระทบต่อสิ่งแวดล้อม** | ◯ | ◯ | ◯ | ◯ | ◯ |

*Skip to question 44*

#### ข้อคิดเห็นเพิ่มเติม

44. มีประเด็นอื่นใดในหัวข้อนี้ที่คุณต้องการแบ่งปันเพิ่มเติมหรือไม่ หรือมีคำถามใดที่เราควรจะถามแต่ยังไม่ได้ถาม

_______________________________________________

_______________________________________________

_______________________________________________

_______________________________________________

_______________________________________________

45. หากท่านยินดีให้เราติดต่อกลับเพื่อสอบถามข้อมูลเพิ่มเติม กรุณาระบุอีเมลด้านล่าง

______________________________

#### 属性情報

ご自身について少しお聞かせください。

46. バイオハッカソンに参加していますか、または過去に参加したことがありますか。（当てはまるものをすべてお選びください）

*Check all that apply.*

- [ ] 2025年 日本バイオハッカソン参加者
- [ ] 過去のバイオハッカソン参加者（国を問わず）
- [ ] バイオハッカソンに参加したことはない

47. 主な専門分野は何ですか。

*Check all that apply.*

- [ ] シーケンス・アナリシス (Sequence analysis)
- [ ] ゲノミクス・トランスクリプトミクス (Genomics and transcriptomics)
- [ ] メタゲノミクス・マイクロバイオーム情報学 (Metagenomics and microbiome informatics)
- [ ] エピゲノミクス・制御ゲノミクス (Epigenomics and regulatory genomics)
- [ ] プロテオミクス・メタボロミクス (Proteomics and metabolomics)
- [ ] 構造バイオインフォマティクス (Structural bioinformatics)
- [ ] システムバイオロジー (Systems biology)
- [ ] 計算進化生物学 (Computational evolutionary biology)
- [ ] ケモインフォマティクス (Cheminformatics)
- [ ] 医療・臨床インフォマティクス (Medical/clinical informatics)
- [ ] 生物統計学 (Biostatistics)
- [ ] 機械学習・AI (Machine learning and AI)
- [ ] データ可視化 (Data visualization)
- [ ] オントロジー構築 (Ontology creation)
- [ ] データ標準化・相互運用性 (Data standards and interoperability)
- [ ] データベース管理 (Database management)
- [ ] HPC・クラウドバイオインフォマティクス (HPC and cloud bioinformatics)
- [ ] フロントエンド開発 (Front-end development)
- [ ] バックエンド開発 (Back-end development)
- [ ] Other: ____________________

48. 所属機関の種類をお選びください。

*Check all that apply.*

- [ ] 教育・研究機関
- [ ] 公的機関
- [ ] 民間企業
- [ ] Other: ____________________

49. 現在どの国で勤務していますか。

____________________

50. 年齢をお知らせください。

*Mark only one oval.*

- 24歳未満
- 25〜34歳
- 35〜44歳
- 45~54歳
- 55歳以上
- 回答しない

51. 性別をお知らせください。

*Mark only one oval.*

- 男性
- 女性
- ノンバイナリー（いずれにも当てはまらない）
- 回答しない
- Other: ____________________

52. 以下の中で、AIの利用状況を最もよく表しているものをお選びください。 *

*Mark only one oval.*

- AI研究者である *Skip to question 54*
- AI開発者である *Skip to question 54*
- 毎日AIを利用している *Skip to question 56*
- 週に数回AIを利用している *Skip to question 56*
- 月に数回AIを利用している *Skip to question 56*
- AIを利用していない *Skip to question 53*
- Other: ____________________

AIを利用していない

53. AIを利用していない理由はありますか。

*Check all that apply.*

- [ ] AIを提供する企業を信頼していない
- [ ] AIという技術は自分の業務に役立たない
- [ ] 費用が高すぎる
- [ ] 結果や基盤となる技術を信頼していない
- [ ] Other: ____________________

*Skip to question 66*

#### AI研究者または開発者

54. AIに関する研究や開発の内容について、簡単にご記入ください。

______________________________
______________________________
______________________________
______________________________
______________________________

55. ご自身の研究や開発の内容に関する論文やコードリポジトリなどを共有頂ける場合はリンクをご記入ください。

______________________________
______________________________
______________________________
______________________________
______________________________

*Skip to question 56*

#### AIの活用分野と影響

56. 最もよく利用しているAIツールについてご記入ください。（複数ある場合は ","で区切ってご記入ください。）

57. AIを利用しているタスクと、タスクにAIを利用する際、以下のうち最も当てはまるものをお選びください。

*Check all that apply.*

| | 編集をほとんど行わず、AIにタスク全体を任せている | AIに下書きを作成させ、その後自分で編集して完成させている | タスクの特定の部分にのみAIを利用している |
|---|---|---|---|
| **コーディング** | ☐ | ☐ | ☐ |
| **研究** | ☐ | ☐ | ☐ |
| **ブレインストーミング** | ☐ | ☐ | ☐ |
| **執筆 / 編集** | ☐ | ☐ | ☐ |
| **教育・教材作成** | ☐ | ☐ | ☐ |
| **翻訳** | ☐ | ☐ | ☐ |
| **個人的な利用（旅行計画、レシピ作成、作曲など）** | ☐ | ☐ | ☐ |

58. 業務やプロジェクトにおいてAIを活用した最大の成功事例についてご記入ください。タスク、使用したAIツール、成果について簡潔にご記入ください。

59. 業務でAIツールを利用する際に直面した課題は何ですか。（当てはまるものをすべてお選びください）

*Check all that apply.*

- [ ] AIツールが複雑で、使いやすくない
- [ ] AIの出力が不正確または信頼できない
- [ ] AIの提案が解釈しづらい
- [ ] 既存のワークフローに統合しにくい
- [ ] 所属機関からの支援が不足している（ソフトウエア、ハードウエア、指導など）
- [ ] 計算コスト / 必要なリソースが高い
- [ ] データの制約（サイズ、品質、アクセス可能性）
- [ ] 倫理的またはプライバシー上の懸念
- [ ] AIへの過度な依存によるミス
- [ ] Other: ____________________

60. AIがうまく機能せず、想定外の問題を引き起こした経験についてご記入ください。そのときに起こったことや、そこから得られた学びについてもご記入ください。

61. AI利用に対する全体的な満足度を1〜5の段階で評価してください。

*Mark only one oval.*

- ( ) 1 - まったく満足していない *Skip to question 62*
- ( ) 2 *Skip to question 62*
- ( ) 3 *Skip to question 62*
- ( ) 4 *Skip to question 63*
- ( ) 5 - 非常に満足している *Skip to question 63*

*Skip to question 62*

#### AIツールの改善に関するご意見

62. AIツールにどのような改善があれば、より満足できると思いますか。

*Check all that apply.*

- [ ] 正確性および信頼性
- [ ] 操作性および効率性
- [ ] 制御およびカスタマイズ性
- [ ] プライバシー・セキュリティおよび倫理
- [ ] Other: ____________________

*Skip to question 63*

#### AI利用に関する所属機関からの支援

63. 所属機関から受けているAI利用に関する支援の程度について、1〜5の段階で評価してください。

*Mark only one oval.*

| | 1 | 2 | 3 | 4 | 5 | |
|---|---|---|---|---|---|---|
| まっ | ( ) | ( ) | ( ) | ( ) | ( ) | 十分に支援されている |

64. 所属機関はどのような支援を提供していますか。（当てはまるものをすべてお選びください）

*Check all that apply.*

- [ ] 独自のAIサービス（オンプレミス / 自社運用）の提供
- [ ] 所属機関が契約するAIソフトウェアやプラットフォームへのアクセス（ライセンス、サブスクリプションなど）
- [ ] ハイパフォーマンスコンピューティング / GPUリソースへのアクセス
- [ ] 専任のAIサポートスタッフやヘルプデスクの設置
- [ ] AIツールやプロンプトエンジニアリングに関する研修・ワークショップの実施
- [ ] 研究ワークフローへのAI統合に関する支援
- [ ] AIの倫理的利用やガバナンスに関する指針の提供
- [ ] AI関連プロジェクトへの資金提供や助成
- [ ] Other: ______________________

*Skip to question 65*

#### AI利用における倫理と責任

65. AIに関連して、以下の課題についてどの程度懸念をお持ちですか。各項目を1〜5の段階で評価してください。

*Mark only one oval per row.*

| | 1 - まったく懸念していない | 2 | 3 | 4 | 5 - 非常に懸念している |
|---|---|---|---|---|---|
| **アルゴリズムにおける偏り** | ⬭ | ⬭ | ⬭ | ⬭ | ⬭ |
| **データのプライバシー・セキュリティ** | ⬭ | ⬭ | ⬭ | ⬭ | ⬭ |
| **知的財産権・所有権** | ⬭ | ⬭ | ⬭ | ⬭ | ⬭ |
| **誤情報/ハルシネーション** | ⬭ | ⬭ | ⬭ | ⬭ | ⬭ |
| **環境への影響** | ⬭ | ⬭ | ⬭ | ⬭ | ⬭ |

*Skip to question 66*

最後に

66. 本調査のテーマに関連して、追加で共有したいことがあればご記入ください。質問項目に含まれていないが触れるべきだと思われる点についてもご記入ください。

67. 今後、追加の質問などでご連絡してもよろしければ、メールアドレスをご記入ください。

This content is neither created nor endorsed by Google.

Google Forms
